# Placode and neural crest origins of congenital deafness in mouse models of Waardenburg-Shah syndrome

**DOI:** 10.1101/2023.10.27.564370

**Authors:** Jamie Tan, Alicia Duron, Henry M. Sucov, Takako Makita

## Abstract

Mutations in the human genes encoding the endothelin ligand-receptor pair *EDN3* and *EDNRB* cause Waardenburg-Shah syndrome (WS4), which includes congenital hearing impairment. The current explanation for auditory dysfunction is a deficiency in migration of neural crest-derived melanocytes to the inner ear. We explored the role of endothelin signaling in auditory development in mice using neural crest-specific and placode-specific *Ednrb* mutation plus related genetic resources. On an outbred strain background, we find a normal representation of melanocytes in hearing-impaired mutant mice. Instead, our results in neural crest-specific *Ednrb* mutant mice implicate a previously unrecognized role for glial support of synapse assembly between auditory neurons and cochlear hair cells. Placode-specific *Ednrb* mutation also caused impaired hearing, resulting from deficient synaptic transmission. Our observations demonstrate the significant influence of genetic modifiers in auditory development, and invoke independent and separable new roles for endothelin signaling in the neural crest and placode lineages to create a functional auditory circuitry.

## Introduction

The auditory system arises from two distinct ectodermal lineages, placode and neural crest. The placode-derived otic vesicle gives rise to the cochlea including mechanosensory hair cells, supporting cells, and the non-melanocyte populations of the stria vascularis. The prosensory domain of the otic placode gives rise to spiral ganglion (auditory) neurons (SGNs); these are bipolar neurons that extend peripheral axons to the hair cells and also project into an ascending auditory pathway (the dorsal and ventral cochlear nuclei) of the central nervous system (Barald and Kelley, 2004; Ladher et al., 2010; Zine and Fritzsch, 2023). SGNs are specified into two functionally distinct types: type I neurons establish afferent synapses on inner hair cells (IHCs), and type II neurons extend afferent axon branches and synapse on outer hair cells (OHCs) (Pyott et al., 2024). The hair cells reside in the cochlear duct, a compartment filled with potassium-enriched endolymph which provides the electrochemical environment that is crucial for hair cell mechanotransduction. The ionic composition (endocochlear potential) of endolymph is maintained by neural crest-derived melanocytes that comprise the intermediate cell layer of the stria vascularis (Steel and Barkway, 1989). Neural crest also gives rise to glia, including satellite cells that enfold individual SGNs in the spiral ganglia, and Schwann cells that are associated with peripheral auditory nerve fibers (D’Amico-Martel and Noden, 1983). In mammals, the final establishment of normal auditory circuitry occurs postnatally (in mice, over the first 2 postnatal weeks). Developmental perturbations that result in defective hair cell mechanotransduction and/or defective mechanosensory circuitry result in congenital sensorineural hearing impairment or deafness, which is a significant human burden (Johnson et al., 2019; Pingault et al., 2010). While the anatomical composition of the normal auditory system is well defined, many of the molecular features that underlie the developmental and maturation events that create auditory functionality or that result in dysfunction are not well studied.

Waardenburg syndrome (WS) is a human congenital disorder characterized by pigmentation abnormalities and sensorineural deafness, and is classified into subtypes based on additional phenotypic components (Gettelfinger and Dahl, 2018). Type 4 Waardenburg syndrome (WS4), also called Waardenburg-Shah syndrome, is distinguished from other WS types because of its association with a congenital gut motility disorder known as Hirschsprung disease (HSCR) (Read and Newton, 1997). Mutations in genes encoding endothelin receptor type B (*EDNRB*) and its ligand endothelin 3 (*EDN3*) are prominently associated with human WS4A and WS4B, respectively (Huang et al., 2022). Importantly, the same mutations in *EDNRB* and *EDN3* are also associated with only HSCR without WS pathologies (HSCR-2 and −3, respectively), indicating variable penetrance of auditory and pigmentation phenotypes.

Defective migration of neural crest cells to a variety of sites is believed to account for all pathological features demonstrated in human WS4. HSCR is characterized by a failure of neural crest-derived enteric nervous system progenitors to migrate into the distal colon, which results in an aganglionic segment in the terminal bowel and thereby a failure of colonic peristaltic contraction. Similarly, melanocytes (pigment cells) throughout the body arise from the neural crest and defective migration of melanoblasts explains pigmentation abnormalities in human WS. In previous studies using inbred mouse strain backgrounds, *Edn3*-*Ednrb* gene mutations recapitulated human WS and HSCR disease pathologies including aganglionic colon and coat color pigmentation defects (Baynash et al., 1994; Hosoda et al., 1994; Ida-Eto et al., 2011; Matsushima et al., 2002). *Ednrb*-deficient mice exhibited an absence of neural crest-derived melanocytes in the stria vascularis (intermediate stria cells) of the cochlea, which like HSCR and coat color alterations could result from a melanocyte migration defect (Deol, 1967; Ida-Eto et al., 2011; Matsushima et al., 2002). These melanocytes generate the endocochlear potential and are essential for hearing, so their absence would explain deafness. To date, absence of neural crest-derived intermediate stria cells is accepted as the primary etiology of sensorineural deafness in WS4.

In this study, we utilized the genetically diverse outbred ICR mouse strain background to examine auditory function in *Edn3*-*Ednrb* global and conditional mutant mice. Unlike prior mouse studies but as observed in humans, all mutant mice exhibited aganglionic colon (HSCR) whereas hearing impairment was variably penetrant. We make two additional key observations. First, hearing impaired mutant mice on this strain background have a normal contribution of melanocytes to the stria vascularis. This challenges the conclusion that deafness in WS is exclusively the result of stria melanocyte migration deficiency; our results instead point to an additional glial-specific (neural crest) role in mechanosensory synapse formation. Second, we show that there is an independent required role for Edn3-Ednrb signaling within the placode-derived sensory neurons to enable auditory functionality. Here, our results implicate an SGN-specific role in synaptic transmission. Our findings establish a dual lineage origin of congenital hearing loss in Edn3-Ednrb type WS4, force reevaluation of the paradigm for auditory defects in the context of these mutations, and demonstrate the complex genetics that cause *Ednrb*-deficiency to manifest in auditory function above or below a hearing threshold.

## Results

### Incomplete penetrance of hearing loss in endothelin signaling mutant mice

Genetic background affects the penetrance and expression of many phenotypes. Variable degrees of pigmentation abnormalities and colonic aganglionosis in *Ednrb* null rodents are examples of this dynamic (Dang et al., 2011). Auditory defects, however, have either been present or absent in these *Ednrb* null rodent models with no indication of variable penetrance (Ida-Eto et al., 2011; Matsushima et al., 2002). We performed click-evoked auditory brain response (ABR) on *Ednrb*- and *Edn3*-null mutant mice that have been maintained in the genetically diverse outbred ICR background. All homozygous mutants for the global *Ednrb* or *Edn3* alleles died within a few days of postnatal day 21 with chronic constipation from colonic aganglionosis (i.e., HSCR) (Poltavski et al., 2024). All animals including control littermates were therefore subjected for ABR at postnatal day 18-19, and their hearing (ABR threshold) determined as the lowest intensity at which a representable ABR waveform could be visually identified (Fig. S1). All control mice (Table S1) showed thresholds in the range of 50-70 (average 60.24±6.91) decibels sound pressure level (dB SPL) (Fig. 1a-b). The hearing threshold of control mice on the ICR background improved to 40-60 (average 49.76±1.02) dB SPL in the next few days (Fig. S2), indicating that hearing is not fully matured at the age of P18-19. In 41 global *Ednrb* mutant mice (from 15 litters generated from 8 independent breeding pairs), 17 mutants showed no response even at the highest sound pressure level (>90dB) and 12 mutants showed an ABR threshold at 80 or 90dB. Surprisingly, 12 mutants showed a comparable response to littermate controls (ABR threshold ≤70dB). These were bilateral traits: all of the first 10 mutants tested as hearing-impaired on one ear exhibited impaired hearing on the other ear, and the first 10 of 10 mutants showing normal hearing on one ear also demonstrated normal hearing on the other ear. Overall, 71% of global *Ednrb* mutant mice exhibited hearing impairment (ABR threshold ≥80dB or no response). Similarly, 61% of global *Edn3* null mice (76 mice from 27 litters generated from 14 independent breeding pairs) showed hearing impairment (19 with no response, 27 with ABR threshold ≥80dB, 30 with ABR threshold ≤70dB). Thus, in this outbred background, auditory defects were very prominent but incompletely penetrant, whereas HSCR was fully penetrant. This resembles the presentation of defects in humans with *EDN3* or *EDNRB* gene mutations (Pingault et al., 2010). Because ICR background mice are albino, pigmentation phenotypes could not be scored.

**Figure 1.**
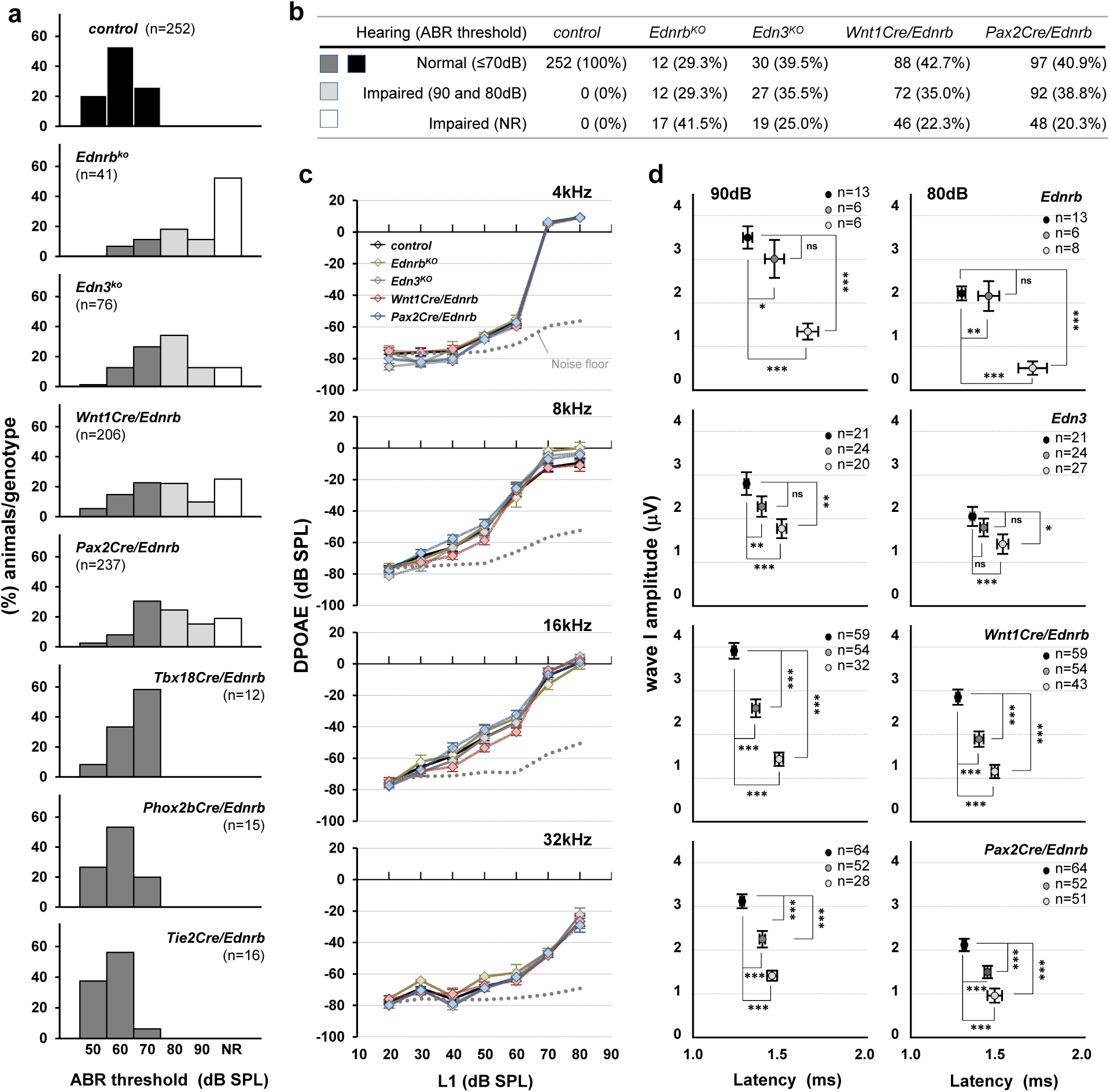
Incomplete penetrance of hearing impairment in Edn3-Ednrb signaling deficient mice. (a) Compiled representation of the percentage of mice of each genotype that exhibited the indicated ABR threshold. Black bars, genetic controls (see also Supplemental Table S1); dark grey bars, hearing mutants; light grey bars, hearing impaired mutants (ABR threshold 80-90dB); white bars, non-hearing mutants (NR=no response at the highest stimulus intensity (90dB)). (b) Table summarizing the number of animals tested and phenotype frequency for global *Ednrb*, global *Edn3*, *Wnt1Cre/Ednrb* and *Pax2Cre/Ednrb* mutant mice. (c) Mean DPOAEs (±SEM) from controls (n=26) and an subset of hearing impaired (ABR threshold≥80dB) *Ednrb* (n=8), *Edn3* (n=15), *Wnt1Cre/Ednrb* (n=15) and *Pax2Cre/Ednrb* (n=13) mutants at L1 intensity level (dB SPL) of the indicated frequency (kHz). L1 is the stimulus level of the first (f1) primary tone (see Methods). (d) Mean amplitude vs latency plots for wave I in ABR waveforms at 90dB (left) and 80dB (right) acquired from hearing mutants (ABR threshold≤70dB, dark grey); hearing impaired mutants (left, ABR threshold 80dB and 90dB, light grey; right, ABR threshold=80dB, light grey) and their littermate controls (black). Error bars; ±SEM. p-values; \**p<0.05*, \*\**p<0.01*, \*\*\**p<0.001*, ns=not significant.

### Edn3-Ednrb signaling is essential not only in neural crest but also in placode lineage for hearing

We defined the sites of Ednrb expression in the cochlea using *Ednrb-eGFP* BAC transgenic mice at P7, a time at which synaptic refinement is just underway shortly before the onset of hearing (P10-13) (Bulankina and Moser, 2012; Johnson et al., 2019). Ednrb expression was detected by GFP immunofluorescence in the lateral wall of the cochlea (stria vascularis) as well as along the entire axis of the spiral ganglia (Fig. 2a). In the spiral ganglia, Ednrb expression was associated with Tuj1^+^ SGNs (Fig. 2b) and Tuj1^+^ (including both afferent and efferent) nerve fibers innervating both IHCs and OHCs (Fig. 2c-2e), and with BLBP^+^ satellite glia (Fig. 2d) and BLBP^+^ and BLBP^−^ Schwann cells (Fig. 2e). In the stria vascularis, Kir4.1 channel-expressing intermediate cells and Kir4.1^−^ basal cells expressed Ednrb-EGFP but not Kir4.1^−^ marginal cells (Fig. 2f). Many but not all epithelial cells of Reissner’s membrane (Fig. 2f) and subsets of PECAM^+^ vascular endothelial cells throughout the cochlea (including ones within Reissner’s membrane; Fig. 2g) also expressed Ednrb. We did not detect Ednrb expression in mechanosensory hair cells (IHCs and OHCs) or supporting cells (Fig. 2c).

**Figure 2.**
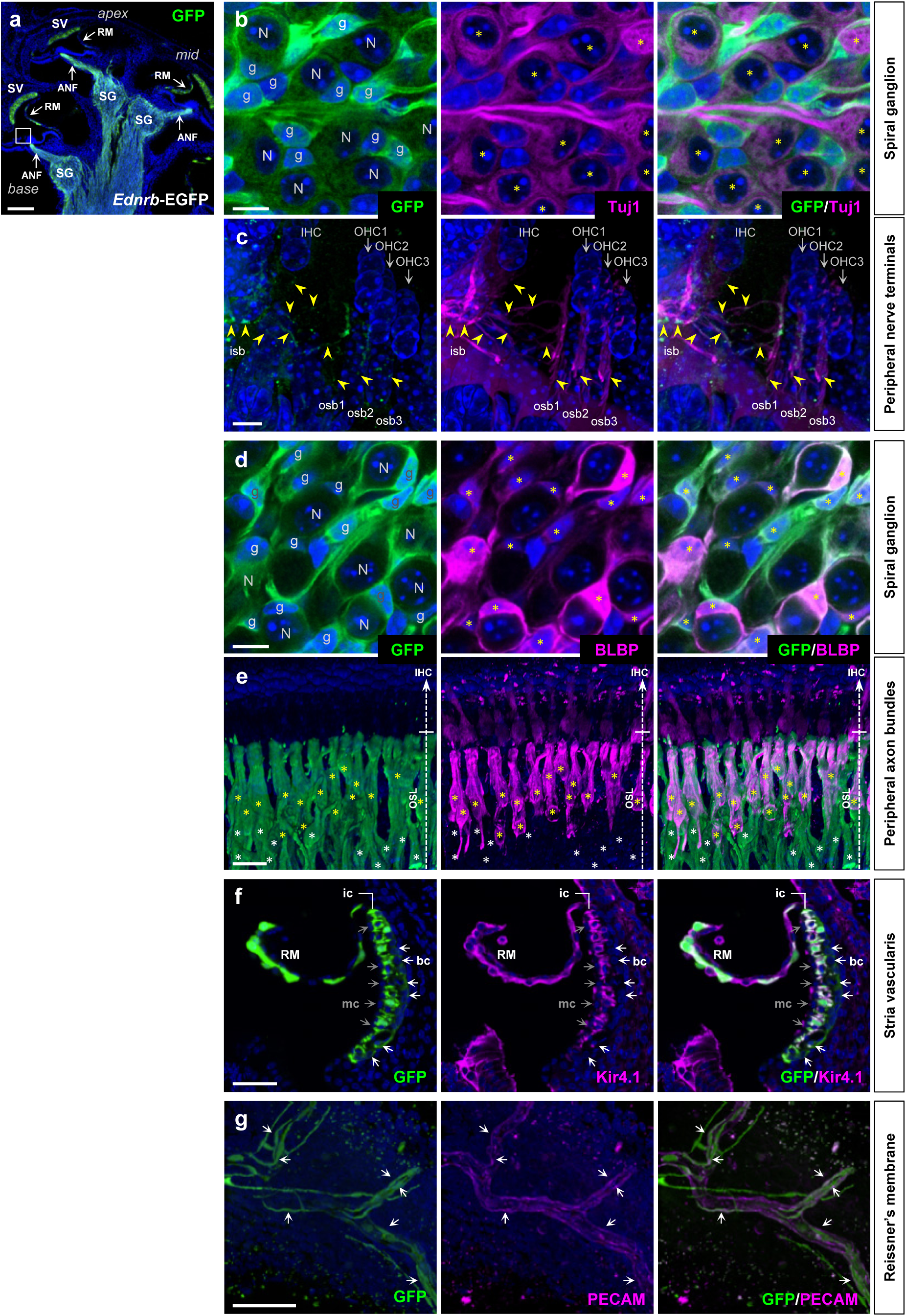
Ednrb expression in P7 cochlea. Sagittal sections of cochlea isolated from P7 *Ednrb-EGFP* BAC transgenic mice stained for GFP only (green; a) or co-stained for neuronal Tuj1 (magenta; b-c), glial BLBP (magenta; d-e), inward rectifier potassium channel Kir4.1 (magenta; f), and vascular endothelial PECAM (magenta; g), and counterstained with DAPI (blue). High magnification views demonstrate Ednrb expression in SGN cell bodies (b) and peripheral nerve terminals (c), in satellite glia in the spiral ganglia (d) and peripheral axon bundles (e), in Kir4.1^+^ intermediate and Kir4.1^−^ basal stria cells (f), and in vascular endothelial cells of the Reissner’s membrane (g, flat mount view). “N” and “g” in (b) and (d) denote spiral ganglion neurons and glia, respectively. Yellow asterisks in (b) denote GFP^+^; Tuj1^+^ spiral ganglion neurons. Yellow arrowheads in (c) point to GFP^+^; Tuj1^+^ axons of spiral ganglion neurons that innervate the inner and outer hair cells corresponding to the boxed area in (a). GFP signal is much stronger in glia than in neurons. Yellow asterisks in (d) denote GFP^+^; BLBP^+^ satellite glia in the spiral ganglion. Among all Schwann cells that are associated with auditory nerve axonal bundles in (e), GFP^+^; BLBP^+^ cells and GFP^+^; BLBP^−^ cells are denoted by yellow and white asterisks, respectively. White and grey arrows in (f) point to GFP^+^; Kir4.1^−^ basal stria cells and GFP^−^; Kir4.1^−^ marginal stria cells, respectively. Arrows in (g) denote GFP^+^; PECAM^+^ vascular endothelial cells of the Reissner’s membrane. Abbreviations: ANF, auditory nerve fibers; bc, basal cell; ic, intermediate cell; IHC, inner hair cell; isb, inner spiral bundle; mc, marginal cell; OHC, outer hair cell; osb, outer spiral bundle; OSL, osseous spiral lamina; RM, Reissner’s membrane; SG, spiral ganglion; SV, stria vascularis. Scale bars: 200μm (a), 10μm (b-d), 25μm (e), 100μm (f), 50μm (g).

Because of this broad expression domain, we evaluated the tissue specific roles of *Ednrb* gene in auditory development. For this purpose, we used *Wnt1Cre* and *Pax2Cre* to drive conditional gene recombination in the neural crest and placode lineages, respectively. When combined with the conditional lineage tracer *R26^nT-nG^* which express nuclear Tomato before and nuclear GFP after Cre-mediated recombination, *Wnt1Cre* activity (visualized by nuclear GFP) was limited to BLBP^+^ satellite cells of the spiral ganglion (Fig. 3a, 3b), BLBP^+^ and BLBP^−^ Schwann cells that are associated with auditory nerve fibers (Fig. 3a, 3c), and Kir4.1^+^ intermediate cells (melanocytes) of the stria vascularis (Fig. 3a, 3d). *Pax2Cre* is active in SGNs (as co-labeled with the pan-neuronal marker HuD) (Fig. 3e, 3f) including Peripherin^+^ type II SGNs (Fig. 3g), and in a variety of cell types in the cochlear duct including hair cells, supporting cells, fibrocytes and epithelium (Fig. 3e). There was no obvious overlap in active domains of *Wnt1Cre* and *Pax2Cre*, although because both are Cre lines they cannot be combined to confirm this point. In spiral ganglia, *Pax2Cre* labeled 94.8±1.1% of SGNs (n=632 from 13 cochlea) and *Wnt1Cre* delineated 95.4±1.0% of satellite cells (n=498 from 11 cochlea). *Pax2Cre* labeled only 75.63±0.63% of hair cells (n=472 from 9 cochlea) (Fig. S3), although because Ednrb is not expressed in hair cells (Fig. 2c), this incomplete recombination efficiency is unlikely to influence this study. Based on the immunolabeling and morphological assessment, *Wnt1Cre* recombination efficiency in the intermediate stria cells was 93.4±1.1% (n=151 from 10 cochlea). Additionally, we used *Tbx18Cre*, *Phox2bCre*, and *Tie2Cre*, which drive high efficiency and specific recombination in otic fibrocytes including basal stria cells (Fig. S4a-b), auditory efferent nerves (Macova et al., 2019) (Fig. S4c-d), and vascular endothelial cells (Fig. S4e-f), respectively. These tissue/cell types express Ednrb (Fig. 2f, 2g) and are relevant for normal hearing (Di Bonito et al., 2023; Furness, 2019; Trune and Nguyen-Huynh, 2012).

**Figure 3.**
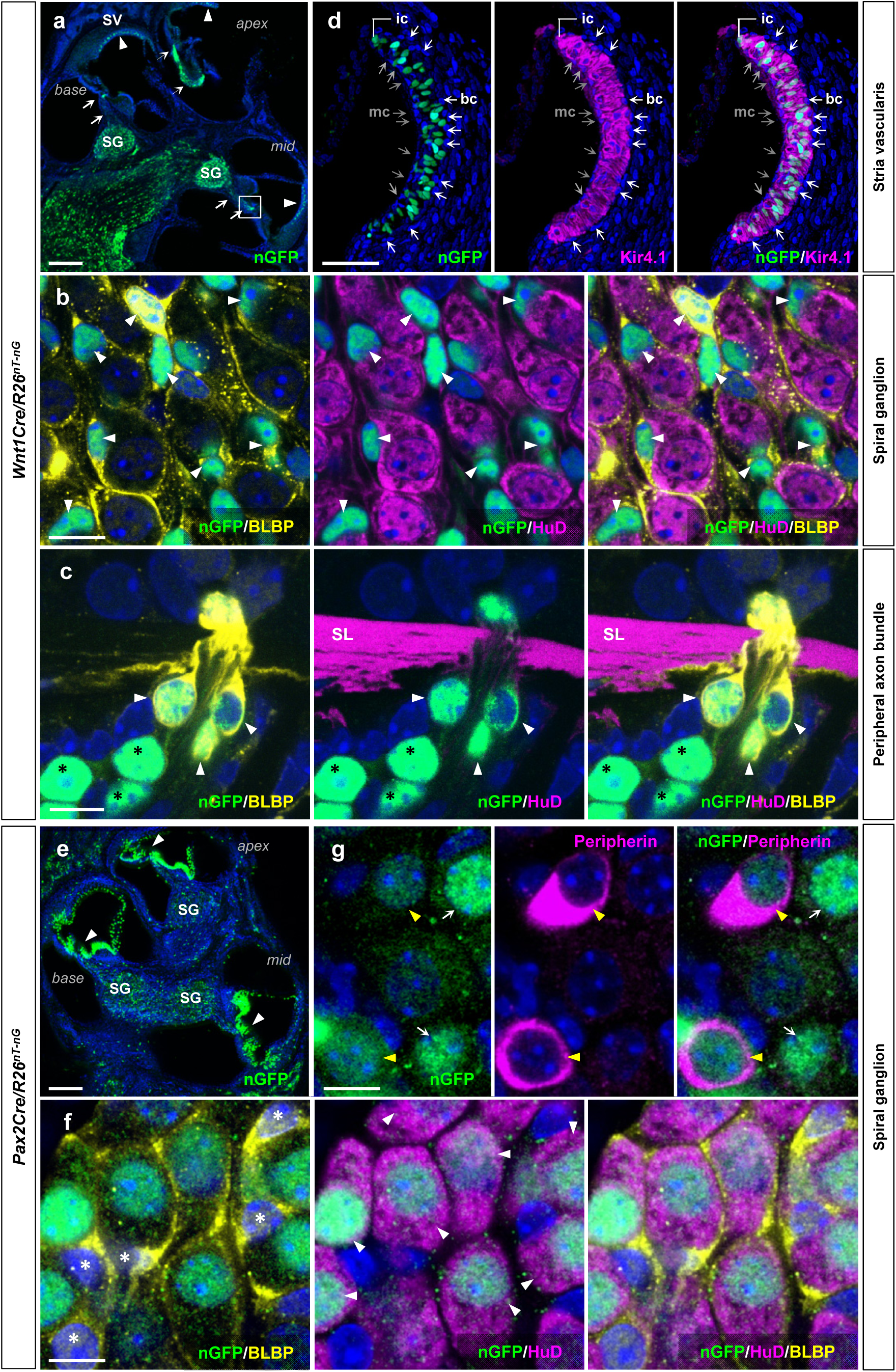
*Wnt1Cre* and *Pax2Cre* delineate distinct cell types in the cochlea. Sagittal sections of cochlea isolated from P7 *Wnt1Cre/R26^nT-nG^*(a-d) and *Pax2Cre/R26^nT-nG^* (e-g) mice stained for GFP (a), costained for neuronal HuD (b-c, f), glial BLBP (b-c, f), Kir4.1 (d), type II neuron-specific Peripherin (g), and counterstained with DAPI (blue). (a) Reporter recombination (nuclear GFP) driven by *Wnt1Cre* occurs in the stria vascularis (arrowheads), and in the spiral ganglion and along their peripheral nerve trajectories (arrows). (b) High magnification view of spiral ganglia in (a) shows *Wnt1Cre* is active in BLBP^+^ satellite cells (arrowheads) but not in HuD^+^ SGNs. (c) A magnified view of the boxed area in (a). *Wnt1Cre* labels BLBP^+^ (arrowheads) and BLBP^−^ (black asterisks) Schwann cells (HuD^−^) along the peripheral nerve bundles. (d) A magnified view of the stria vascularis in (a) shows the activity of *Wnt1Cre* in Kir4.1^+^ intermediate stria cells, but not in Kir4.1^−^ basal (white arrows) and medial (grey arrows) stria cells. (e) Reporter recombination by Pax2Cre occurs in vast majority of cochlear epithelium (arrowheads) and in the spiral ganglion. (f-g) High magnification views of spiral ganglia in (e) demonstrate that *Pax2Cre* is exclusively active in HuD^+^ SGNs (f; white arrows) including Peripherin^+^ type II neurons (g; yellow arrows) but not in BLBP^+^ satellite cells (f; asterisks). Abbreviations: SL, spiral lamina. Scale bars; 200μm (a, e), 10μm (b-c, f-g), 100μm (d).

We crossed each of these Cre drivers with the conditional *Ednrb* allele and performed ABR evaluation. As we recently described (Poltavski et al., 2024), the enteric nervous system has independent requirements for the neural crest and placode lineages such that *Wnt1Cre/Ednrb* and *Pax2Cre/Ednrb* mutants both die around P21 with Hirschsprung disease. Thus, ABR analysis on conditional *Ednrb* mice was conducted at P19, just as done with global mutant mice described above. The analysis included 206 *Wnt1Cre/Ednrb* mutants (92 litters from 39 breeding pairs), 237 *Pax2Cre/Ednrb* mutants (103 litters from 36 breeding pairs), 12 *Tbx18Cre/Ednrb* mutants (4 litters from 4 breeding pairs), 15 *Phox2bCre/Ednrb* mutants (9 litters from 5 breeding pairs) and 16 *Tie2Cre/Ednrb* mutants (4 litters from 2 breeding pairs)(Fig. 1a). Controls included Cre-positive and *Ednrb* heterozygous mice from these same litters (Table S1). Hearing impairment was evident in both *Wnt1Cre/Ednrb* and *Pax2Cre/Ednrb* with incomplete penetrance (57% and 59%, respectively), just as observed in global *Ednrb* and *Edn3* mutant mice. Thus, *Ednrb* gene function is independently required in the neural crest and placode lineages for auditory function. All *Tbx18Cre/Ednrb*, *Phox2bCre/Ednrb* and *Tie2Cre/Ednrb* mutants exhibited ABR thresholds comparable to controls (Fig. 1a-1b), indicating a lack of *Ednrb* requirement in these lineages.

To further define hearing impairment observed in these mice, we tested a subset of global *Ednrb* (n=8), global *Edn3* (n=14), *Wnt1Cre/Ednrb* (n=15) and *Pax2Cre/Ednrb* (n=13) mutants that were ABR non-responsive at 90dB for Distortion Product Otoacoustic Emissions (DPOAEs) at P20. All of these deaf mice presented a comparable DPOAE response to control mice (n=26) in the stimulus range of 4-32 kHz, indicating that cochlear amplifier function is fully intact in all four mutant backgrounds (Fig. 1c).

We further evaluated the ABR waveforms of hearing (ABR threshold ≤70dB) and hearing impaired (ABR threshold 80 or 90dB) mutants in each of these four mutant backgrounds. Wave I represents the activity of primary afferent neurons (SGNs), and its amplitude reflects the stimulus intensity and the number of mechanosensory axons that are simultaneously activated, whereas its latency reflects the duration between the initial stimulus and the peak of wave. Hearing impaired mutants of all four genotypes exhibited significantly lower wave I amplitude and longer latency compared to their controls at the stimulus intensity range (90dB and 80dB) at which they responded (Fig. 1d, Fig. S1). This indicates activation of fewer SGNs and less synchronicity in SGN activation upon stimulation.

Interestingly, although not as severely compromised as in hearing impaired mutants, hearing mutants (i.e., mutants with ABR threshold ≤70dB) also displayed less amplitude and longer latency relative to controls; this was statistically significant in *Wnt1Cre/Ednrb* and *Pax2Cre/Ednrb* mutants because of greater sample number but also showed the same trend in global mutants (Fig. 1d). The high specificity and recombination efficiency of *Pax2Cre* and *Wnt1Cre* imply that variable recombination does not explain phenotypic range and penetrance. Rather, we suggest that additional genetic factors (phenotypic modifiers) in the diverse genetic background of our outbred colony converge with Edn3-Ednrb signaling to determine whether hearing is normal, impaired, or fully absent (see Discussion).

### Melanocyte deficiency is not associated with hearing impairment in *Ednrb*-deficient mice

Previous mouse studies with global *Ednrb* mutants in inbred strain background reported a complete absence of intermediate cells in the stria vascularis (Ida-Eto et al., 2011; Matsushima et al., 2002), and thus the etiology of hearing loss in WS4 is thought to result from impaired migration of neural crest-derived melanocytes to the stria vascularis. To address the presence or absence of cochlear melanocytes in *Wnt1Cre/Ednrb* mutant mice, we crossed the *R26^td-Tomato^* reporter into the *Wnt1Cre/Ednrb* background and visualized neural crest-derived cells at P19 in the cochlea of 6 hearing mutants and 15 hearing impaired mutants generated from 3 independent breeding pairs (Fig. 4). Surprisingly, in both hearing (ABR threshold ≤70dB) and hearing impaired mutants, *Wnt1Cre*-labeled cells were present in the intermediate layer of the stria vascularis. Moreover, these cells were functionally mature as evidenced by expression of the inwardly rectifying potassium channel Kir4.1. In several mutants we observed various mild anomalies including a partial loss of *Wnt1Cre*-labeled cells (Fig. 4c, 4e, 4g), a partial loss of Kir4.1 expression in *Wnt1Cre*-labled intermediate stria cells (Fig. 4c, 4e, 4g), and misexpression of Kir4.1 in non-Wnt1Cre-labeled marginal cells (Fig. 4c, 4g). However, these anomalies were also present in hearing mutants (Fig. 4g). Significantly, the majority of hearing impaired mutants (and of hearing mutants) had a fully normal organization of the stria vascularis. Although very few were analyzed in this manner, we also observed normal population and maturation of intermediate stria cells in global *Ednrb* deaf mutants (Fig. 4f, 4g). Thus, in the ICR background of this colony, cochlear melanocyte defects in *Ednrb* mutants are variable and at most are limited in scale. More importantly, in these mice melanoblast migration or melanocyte deficiency cannot be the primary cause of hearing loss.

**Figure 4.**
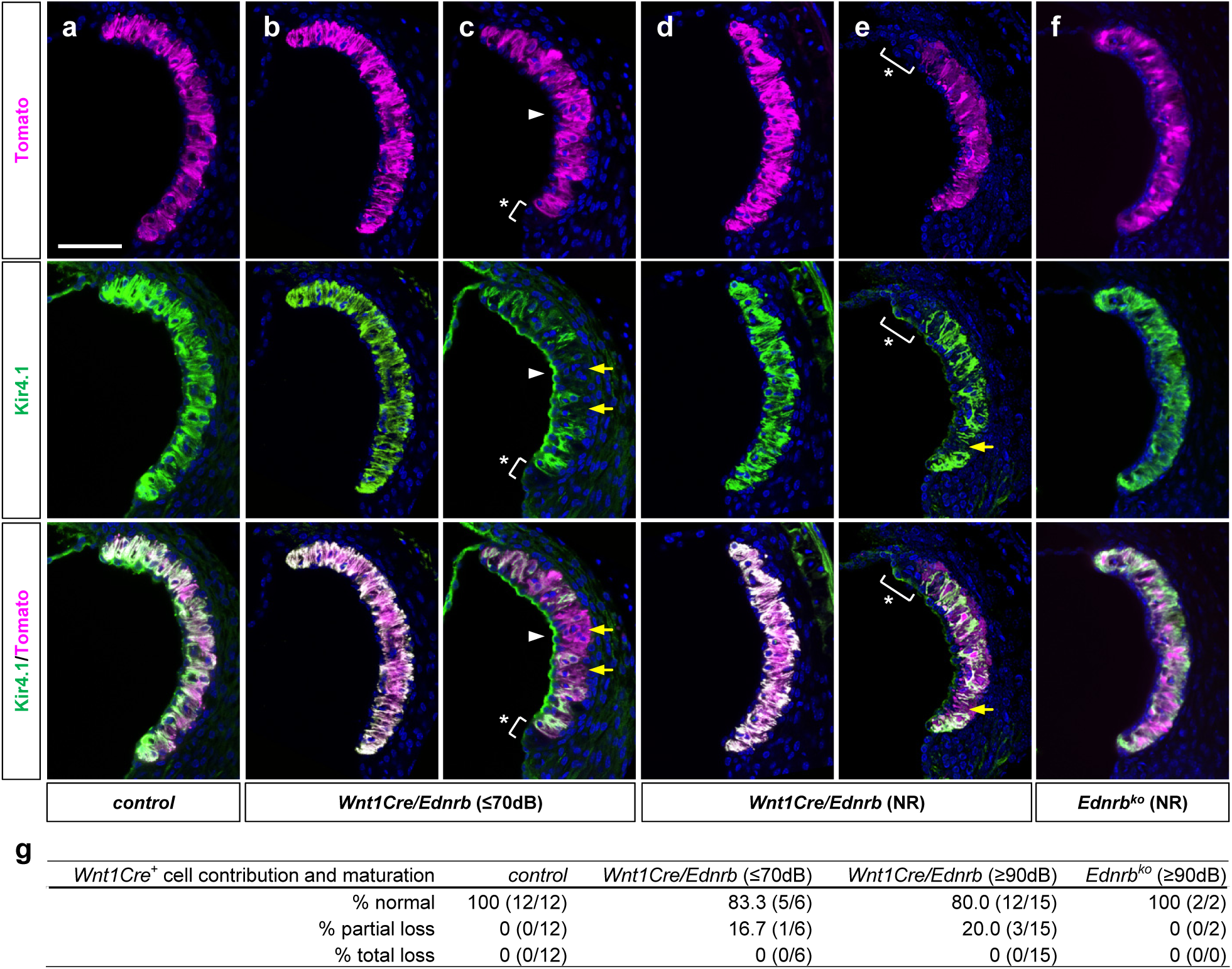
Contribution of *Wnt1Cre*-labeled melanocytes to the Kir4.1^+^ intermediate stria cells in hearing and hearing impaired *Ednrb*-deficient mice. Confocal (z-stack) images of the stria vascularis of P19 hearing (ABR threshold≤70dB) *Wnt1Cre/Ednrb/R26^td-Tomato^*mutant (b-c), hearing impaired (ABR threshold=NR) *Wnt1Cre/Ednrb/R26^td-Tomato^* mutants (d-e), a littermate control (a), and hearing impaired (ABR threshold=NR) global *Ednrb* mutant crossed into the *Wnt1Cre/R26^td-Tomato^*background (f) immunostained for Kir4.1 (green) and counterstained with DAPI (blue). (c, e) Asterisks denote the portion of the stria vascularis that does not contain *Wnt1Cre* lineage derived Kir4.1^+^ intermediate cells. Arrows point out *Wnt1Cre* lineage^+^ cells that are lacking Kir4.1 expression. Arrowheads point out misexpression or mislocalization of Kir4.1 at the apical surface of the marginal stria cells. Scale bar: 100μm. (g) Table summarizing the number of animals tested and the phenotype (intermediate cell number and maturation) frequency for *Wnt1Cre/Ednrb* and global *Ednrb* mutant mice.

### Contribution of glial *Ednrb*-deficiency to defective mechanosensory synapse formation

If melanocyte deficiency in the stria vascularis does not explain hearing impairment, we asked if hearing loss instead arises from auditory nervous system defects. We performed immunofluorescence analysis for Tuj1, synaptophysin (SYP), and C-terminal-binding protein 2 (CtBP2) in *Wnt1Cre/Ednrb* mutant cochlea to evaluate auditory nerve fibers, auditory synapses, and synaptic ribbon profiles. In wholemount (flat mount) views of control cochlea at P19, Tuj1^+^ nerve endings have assembled postsynaptic SYP clusters and presynaptic CtBP2 at each IHC and OHC (Fig. 5a). Sagittal views confirmed that presynaptic CtBP2^+^ ribbons were assembled towards the basolateral surface of the IHCs and OHCs which juxtaposed with post synaptic SYP (Fig. S5a-c). Whereas hearing mutants displayed almost normal pre- and postsynaptic assembly (Fig. 5b, 5d-g: grey), in hearing impaired mutants there was notable disorganization of SYP clusters as well as CtBP2^+^ ribbons on both IHCs and OHCs (Fig. 5c, 5d-g: red). This was particularly prominent at OHCs (Fig. 5g), but also observed at IHCs (Fig. 5e). The typical characteristic of this disorganization was poor alignment of postsynaptic SYP and presynaptic CtBP2, as shown by inconsistent size/volume of SYP clusters and more anteriorly distributed CtBP2 ribbons in both inner and outer hair cells (Fig. S5d-f). Synaptic ribbons facilitate rapid, precise and continuous neurotransmission and are critical for auditory perception. Postsynaptic SYP showed a corresponding pattern, being more altered in hearing impaired OHCs (Fig. 5f) than IHCs (Fig. 5d). As SYP staining represents both efferent and afferent synapses, the disruption of afferent SYP^+^ synapses at both IHCs and OHCs in hearing impaired *Wnt1Cre/Ednrb* mutants is likely much greater than what is presented and quantified in Fig. 5. These results imply that defective mechanosensory synapse formation is the primary phenotype that accounts for auditory impairment in *Wnt1Cre/Ednrb* mutants. Without proper afferent synaptic organization, auditory function would be compromised or eliminated.

**Figure 5.**
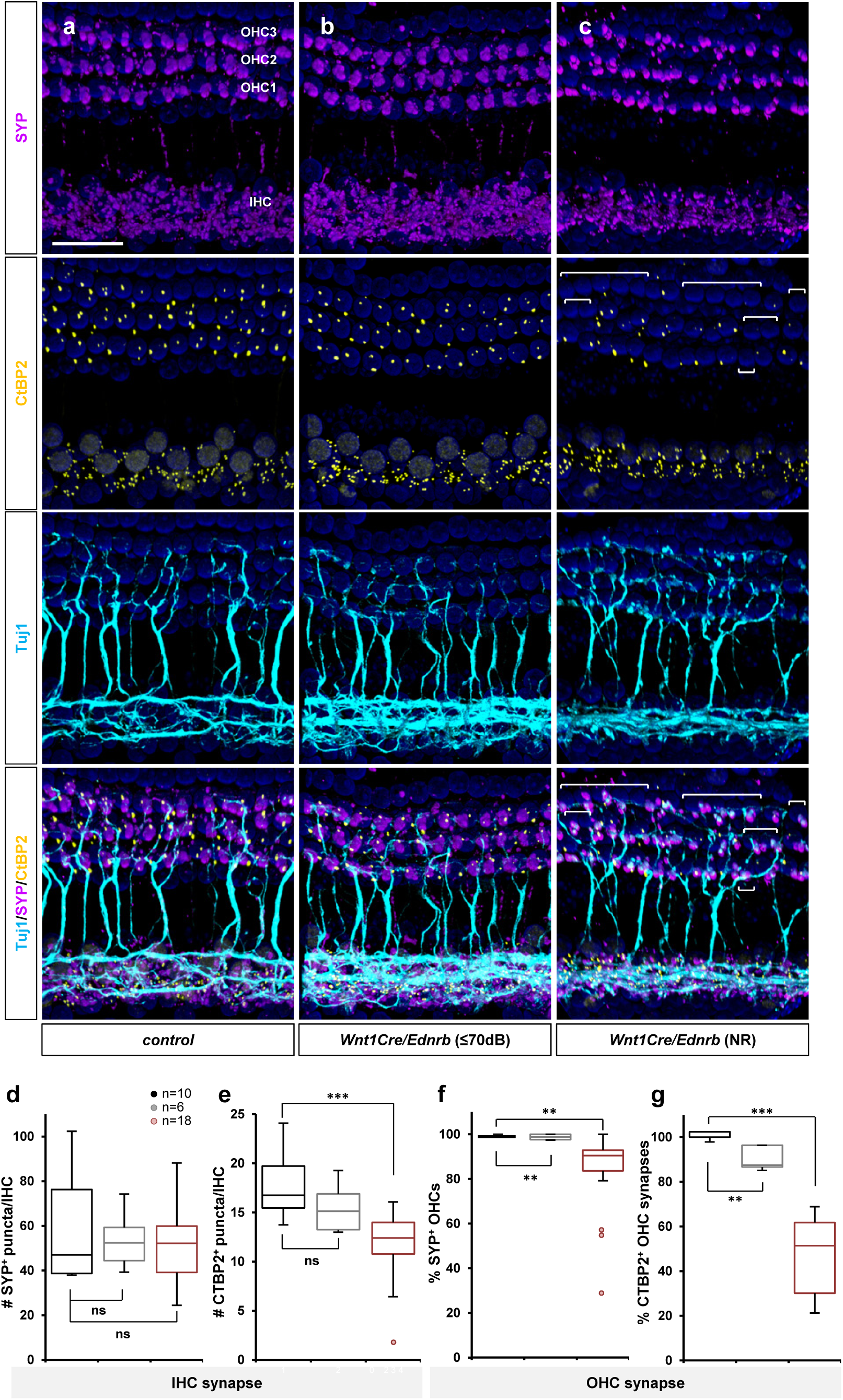
Defective synapse formation in hearing impaired *Wnt1Cre/Ednrb* mutant mice. Whole-mount preparations of cochlea isolated from P19 *Wnt1Cre/Ednrb* mutant mice (b-c) and a littermate control (a) immunostained for postsynaptic SYP (magenta), presynaptic CtBP2 (yellow), neuronal Tuj1 (cyan), and counterstained with DAPI (blue). The image shown in (c) is from a hearing impaired *Wnt1Cre/Ednrb* mutant of median synaptic phenotype. OHCs in the bracketed areas lack CtBP2^+^ synaptic ribbons. Scale bar: 50μm. (d-e) Box and whisker plots for the number of SYP^+^ (d) and CtBP2^+^ (e) puncta per IHC in control (black), hearing *Wnt1Cre/Ednrb* mutant (grey), and hearing impaired *Wnt1Cre/Ednrb* mutant (ABR threshold ≥80dB; red) mice. (f-g) Box and whisker plots for the percentage of OHCs with postsynaptic SYP^+^ staining (f) and presynaptic CtBP2^+^ staining (g) in control (black), hearing *Wnt1Cre/Ednrb* mutant (grey), and hearing impaired *Wnt1Cre/Ednrb* mutant (ABR threshold ≥80dB; red) mice. Extremely compromised outliers in (e-f) were plotted individually and not included in the statistical analysis. p-values: **p<0.01, ***p<0.001, ns=not significant.

*Wnt1Cre* is not active in SGNs nor in their primary afferent targets (IHCs and OHCs), all of which are placode-derived. *Wnt1Cre* is active in the mid/hindbrain region which includes the olivocochlear nuclei, from which efferent nerves originate. For two reasons this source is unlikely to contribute to auditory impairment in *Wnt1Cre/Ednrb* mutants. First, the efferent system is primarily involved in auditory discrimination and cellular protective roles rather than in primary auditory perception (Ashmore et al., 2023; Elgoyhen and Katz, 2012) and so is outside the realm of ABR measurement. More decisively, *Ednrb* conditional mutation with efferent nerve-specific *Phox2bCre* did not result in hearing impairment (Fig. 1a). The one remaining *Wnt1Cre* sublineage that could plausibly account for defective synapse formation is glia. *Wnt1Cre*-labeled glial cells are abundantly present in the cochlea, as satellite cells tightly associated with SGNs (Fig. 3b) and as Schwann cells associated with afferent nerve fibers (Fig. 3c). Satellite cells and Schwann cells both express Ednrb (Fig. 2). Glial cells play important roles in many aspects of neural development and maturation, including synapse formation in both central and peripheral nervous system including auditory afferent innervation (Hanani and Verkhratsky, 2021; Mao et al., 2014). Anatomically, there was no obvious disruption in the number or distribution of glial cells in the spiral ganglia of mutant mice (Fig. S6). This suggests a functional deficiency in glial cells as the underlying condition that results in hearing impairment. Because synaptic defects observed here are between SGNs and hair cells, both of which are placode-derived, these results imply that an initial endothelin signaling process in glia, acting through the afferent axons of SGNs, remotely supports mechanosensory synapse organization at hair cells, and which is compromised in hearing impaired *Wnt1Cre/Ednrb* mutants.

### An intrinsic requirement for Ednrb in type I SGN activation/excitation

Unlike *Wnt1Cre/Ednrb* mutants, hearing impaired *Pax2Cre/Ednrb* mutant mice displayed normal presynaptic CtBP2 and postsynaptic SYP profiles at both IHCs and OHCs (Fig. S7a-c). This indicates that the underlying cell and molecular events that impact on auditory perception in *Pax2Cre/Ednrb* mutants are different from the synapse defect observed in *Wnt1Cre/Ednrb* mutant mice. *Pax2Cre* is active in the majority of otic cell types (Fig. 3e), although Ednrb is not expressed in most of these that are relevant for auditory function (e.g., hair cells, supporting cells, etc.). *Pax2Cre* denotes subsets of Ednrb-expressing otic fibrocytes, but the normal hearing of *Tbx18Cre/Ednrb* mutants (Fig. 1) refutes basal stria cells (Fig. S4b) as the cause of hearing impairment. SGNs are the main cell type that is within the *Pax2Cre* recombination domain and expresses Ednrb, suggesting an intrinsic requirement for Ednrb function in SGNs. Although postsynaptic SYP labeling at sensory hair cells was normal (Fig. S7), we noted a significant reduction in vesicular SYP staining in *Pax2Cre/Ednrb* mutant SGNs (Fig. 6c). This phenotype was observed in all hearing impaired *Pax2Cre/Ednrb* mutants analyzed, whereas SGNs of hearing *Pax2Cre/Ednrb* mutants showed comparable levels of vesicular SYP^+^ area to control mice (Fig. 6a-6c, 6g). Vesicular transport in neurons is a reflection of synaptic transmission. Thus, these observations suggest that *Ednrb* deficiency in SGNs results in impaired neural activation.

**Figure 6.**
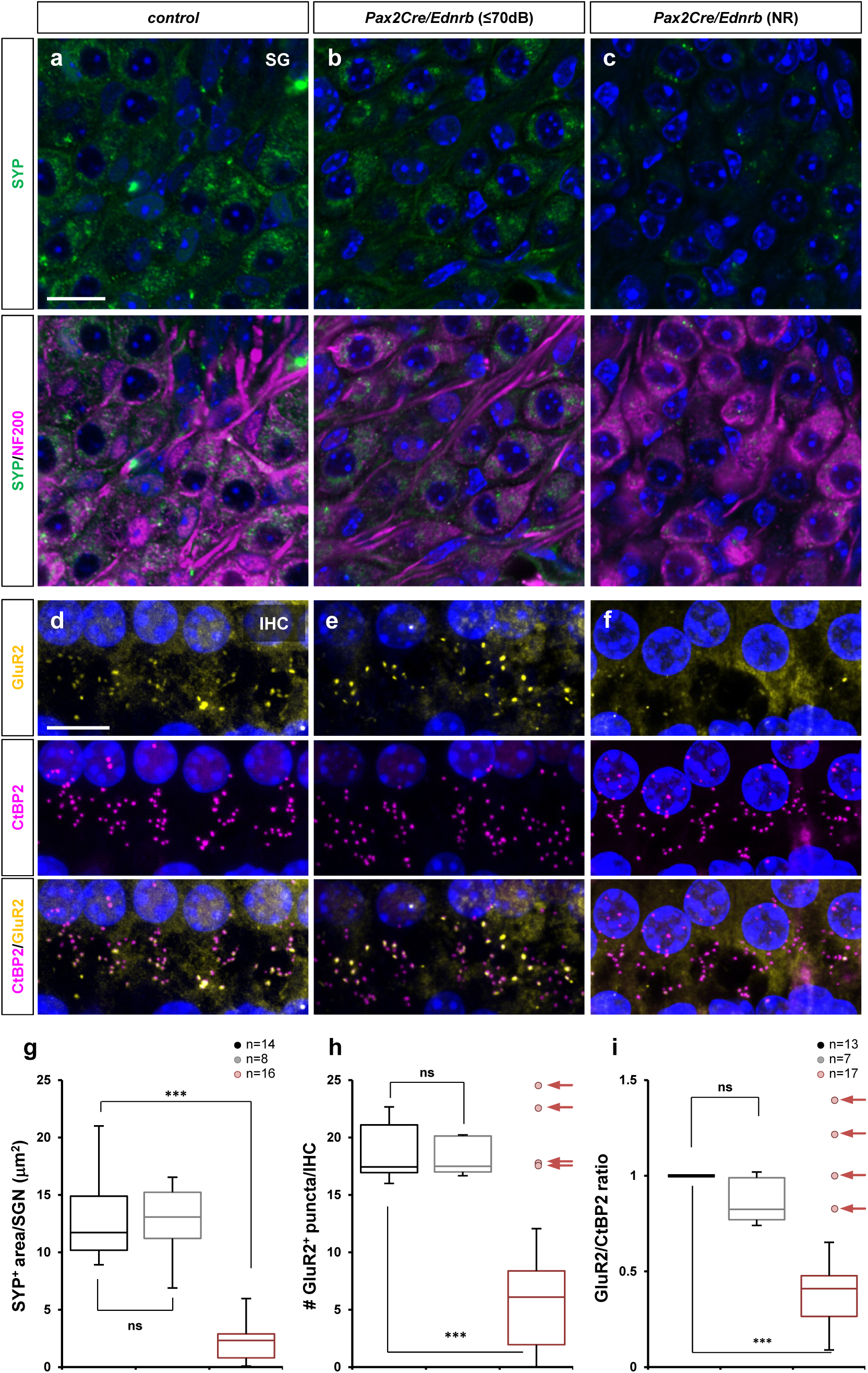
Defective SGN activation in hearing impaired *Pax2Cre/Ednrb* mutant mice. (a-c) High magnification views of P19 spiral ganglion sections from hearing *Pax2Cre/Ednrb* (b), hearing impaired *Pax2Cre/Ednrb* (c) mutant mice and a littermate control (a) immunostained for SYP (green), and NF200 (magenta), and counterstained with DAPI (blue). Compiled representation of SYP^+^ area per SGN in each group is shown in g (see Methods). (d-f) Whole-mount preparation of cochlea isolated from P19 hearing *Pax2Cre/Ednrb* (e), hearing impaired *Pax2Cre/Ednrb* (f) mutants and a littermate control (d) immunostained for GluR2 (yellow), and CtBP2 (magenta), and counterstained with DAPI (blue). Compiled representation of GluR2^+^ puncta number per IHC and GluR2/CtBP2 ratio per IHC are shown in h and i, respectively (see Methods). Scale bars: 10μm. (g) Box and whisker plots for the SYP^+^ area per SGN in control (black), hearing *Pax2Cre/Ednrb* mutant (grey) and hearing impaired mutant (ABR threshold=NR; red) mice. (h-i) Box and whisker plots for the number of GluR2^+^ puncta per IHC (h) and the ratio of GluR2/CtBP2 puncta (i) in control (black), hearing *Pax2Cre/Ednrb* mutant (grey) and hearing impaired mutant (ABR threshold=NR; red) mice. Arrows in (h) and (i) represent hearing impaired *Pax2Cre/Ednrb* mutants exhibiting a normal GluR2 profile at the IHC synapse that were not included in the statistical analysis. p-values: **p<0.01, ***p<0.001, ns=not significant.

Glutamate is the primary neurotransmitter released by hair cells to activate SGNs, which receive this signal via AMPA receptors (Reijntjes and Pyott, 2016). To test if impaired glutamate signaling accounts for reduced SGN activity in hearing impaired *Pax2Cre/Ednrb* mutant mice, we examined the expression and localization of the AMPAR subunit GluR2 and presynaptic CtBP2. We screened a total of 17 hearing impaired *Pax2Cre/Ednrb* mutants and 7 hearing *Pax2Cre/Ednrb* mutants. In hearing *Pax2Cre/Ednrb* mutants, GluR2^+^ clusters were detected at IHCs at a comparable level to control IHCs (Fig. 6e, 6h), and each GluR2^+^ cluster was located in conjunction with a presynaptic CtBP2^+^ puncta (Fig. 6e, 6i), as also observed at IHCs of control mice (Fig. 6d, 6h-6i). However, in hearing impaired *Pax2Cre/Ednrb* mutant mice, the number of GluR2^+^ clusters on average was significantly reduced, although those GluR2^+^ clusters that were present at IHCs were properly juxtaposed with CtBP2 (Fig. 6f, 6h-6i). Interestingly, this GluR2 abnormality was incompletely penetrant: it was found in 13 out of 17 hearing impaired *Pax2Cre/Ednrb* mutants (Fig. 6h-6i), but a subset of 4 hearing impaired mutants (Fig. 6h-6i; arrows) exhibited a normal number of GluR2^+^ clusters even as they still presented an impaired vesicular SYP^+^ profile (Fig. 6c, 6g). This divergence is unlikely to simply be measurement error but more likely reflects phenotypic variation associated with the outbred background of our colony. In sum, in hearing impaired *Pax2Cre/Ednrb* mutants, alteration of vesicular SYP and reduced GluR2 localization together lead to the conclusion that the primary lesion is an SGN-autonomous defect in synaptic transmission. As also for *Wnt1Cre/Ednrb* mutants, the variable penetrance of hearing phenotypes is inferred to reflect the activity of unlinked genetic alleles (modifiers) that are variable between individual mice in this study.

## Discussion

Congenital deafness is among the more prevalent chronic conditions seen in human infants. Genetic causes are thought to explain the majority of cases in developed countries. In many cases the genetic basis of deafness is unknown, and even where causative gene mutations are known, the explanation for how these are transduced into auditory development or function has often been unclear. In this analysis of *Edn3* and *Ednrb* mutant mice, we reach several new insights that have not been appreciated in prior studies. First, hearing impairment is a variable trait in endothelin signaling mutant mice as observed in human. Second, a deficiency of melanocyte migration to the stria vascularis does not explain deafness in this WS4 model. Lastly, the sites of action of endothelin signaling that account for WS4 hearing loss include not only neural crest-derived but also placode-derived cells in the cochlea, and their developmental and mechanistic roles are clearly distinct in each embryonic cell lineage. To a significant extent, these insights were possible to recognize because of the outbred diverse genetic background of our colony. In addition, we could distinguish these processes through tissue-specific conditional analysis that has not previously been conducted.

The paradigm that melanocyte migration deficiency explains deafness in Waardenburg syndromes and in particular in WS4 is strongly entrenched. This was reasonable: Hirschsprung disease represents migration failure of enteric progenitors to the colon, pigmentation defects represent failure of melanocyte migration into the skin, and the intermediate cells of the stria vascularis are melanocytes that derive from neural crest and must migrate to become incorporated into the inner ear. In mouse development, melanoblasts appear in the future lateral wall of the cochlear duct as early as embryonic day 12.5 and complete migration to the intermediate layer of the stria vascularis by postnatal day 0 (Bonnamour et al., 2022; Renauld et al., 2022). It is worth noting that numerous other neural crest lineages that also require migration do so normally in *Edn3* and *Ednrb* mutants and in other WS models (e.g., dorsal root and sympathetic chain ganglia, the outflow tract of the heart, adrenal medulla, etc.), so impaired migration of stria intermediate cells in *Edn3* or *Ednrb* mutants is not a foregone conclusion. Nonetheless, several past studies using inbred *Ednrb* rodent models (Ida-Eto et al., 2011; Matsushima et al., 2002) (*Edn3* has not been studied in this regard) have concluded defective migration of cochlear melanocytes as evidenced by total absence of intermediate stria cells. We do not see evidence of this in our mice. One possibility is that melanocyte migration is better supported in the strain background (ICR) of our mice. Regardless, our results indicate that endothelin signaling has multiple roles in the neural crest cell lineage of the inner ear, including in auditory synaptic assembly as observed here. This latter role has not been previously recognized.

We note that our assessment of mature melanocytes in the stria vascularis is based on the visible presence of neural crest lineage-labeled cells expressing Kir4.1, but this is not a functional assessment of endocochlear potential. It has been reported in a variety of mouse models of hearing loss that DPOAE reflects endocochlear potential (Ingham et al., 2021; Tian et al., 2021; Xia et al., 2007): animals with reduced DPOAEs were associated with reduced endocochlear potential, whereas animals with normal DPOAEs showed normal endocochlear potential. Thus, together with the normal presence of Kir4.1-expressing intermediate stria cells, we infer that endocochlear potentials in our hearing impaired *Wnt1Cre/Ednrb* mutants are normal enough to generate normal DPOAEs (Fig. 1c). We have not yet ruled out if *Ednrb*-deficient intermediate cells influence the adjacent marginal cell functions in homeostatic regulation of endolymph Ca^2+^ and HCO_3_ ^−^ (and other cations/anions) that could also influence hearing threshold (Tian et al., 2021; Wangemann et al., 2004; Wangemann et al., 2007).

Alternatively, our studies implicate glial cells as a candidate site of Ednrb activity to explain deafness in *Wnt1Cre/Ednrb* mutants. To a certain extent, our interest in glia is based on exclusion of other candidate neural crest cell types (melanocytes and cranial efferent nerves). Formal proof of a glial-specific Ednrb role will require an appropriate glia-specific Cre line which we do not yet have. Nonetheless, such a role is quite reasonable: for example, in mice with conditional deletion of *Sox10* using *Wnt1Cre*, satellite/Schwann cells were absent in the spiral ganglia by E16.5, and in their absence, placode-derived SGNs failed to migrate to appropriate sites and also failed to project their afferent axons to appropriate targets (Mao et al., 2014; Szeto et al., 2022). Apparently, glial cells play crucial roles in guiding cochlear neuron migration and axonal growth during embryonic development at least by physical contact (and possibly by molecular interactions). In our *Wnt1Cre/Ednrb* mutants, along with melanocyte migration, cochlear glia migrate normally and form spiral ganglia with placode-derived SGNs at the appropriate location with no obvious defects in their size and morphology. Furthermore, *Ednrb*-deficient satellite cells normally express Kir4.1 and are capable of enfolding individual SGN (Fig. S6c). We propose that glial endothelin signaling is dispensable in the embryonic phase of cochlear development but crucial for functional maturation of auditory circuitry: in response to Edn3 and mediated through Ednrb, glial cells remotely promote afferent synapse formation either by secreting paracrine factors that act on SGNs or through cell contact-mediated activation/suppression of relevant signaling pathways in SGNs. Additional high-resolution analysis such as transmission electron microscopy may be warranted to formally validate defective synapse formation in hearing impaired *Wnt1Cre/Ednrb* mutants. As presented by the variable severity of deafness in hearing impaired *Wnt1Cre/Ednrb* mutants (Fig. 1a-b, 1d), we predict that primary *Ednrb*-deficiency makes spiral ganglion glia vulnerable to additional and/or epistatic effects of other (modifier) genes, and which ultimately manifest impairment in synapse formation. For a better understanding of the mechanistic basis of glial Ednrb-mediated hearing loss, it will be informative to identify those modifier genes and to define how these genes independently interact with primary *Ednrb*-deficiency in spiral ganglion glia.

Our studies have also revealed a placode-specific role for endothelin signaling in auditory functionality. A hint of this role appeared in one prior study in which transgenic reexpression of Ednrb in SGNs of global *Ednrb* mutant mice resulted in partial improvement in hearing (Ida-Eto et al., 2011). This was associated with rescue of SGN neurodegeneration. We did not observe SGN neurodegeneration in our global and conditional mutant mice, which is possibly because neural survival is better supported in the ICR background or because the postnatal day 19 time point that we use for our study (due to HSCR lethality) is not old enough to detect neurodegeneration. We also did not observe any alteration in spiral ganglion size and morphology including number and distribution of Peripherin^+^ type II neurons as previously described (Elliott et al., 2021). Instead, our data implicate SGN synaptic transmission as the primary lesion in *Pax2Cre/Ednrb* mutants. The one common feature of all hearing impaired *Pax2Cre/Ednrb* mutants was a reduction of SYP^+^ vesicles, which is an indication of diminished synaptic activity. We observed diminished synaptic GluR2 in most but not all deaf mutants; this is a likely contributor to diminished vesicular transport in at least these mice. One possibility is that Ednrb regulates GluR2 expression or localization, although if so there are clearly additional mechanisms involved that account for normal GluR2 in hearing *Pax2Cre/Ednrb* mutants and as well in a small subset of deaf mutants.

We believe that genetic modifiers in the outbred ICR strain background of our colony account for the variable penetrance of hearing impairment in global and conditional mutant backgrounds. Strain background can also explain variability in GluR2 localization in *Pax2Cre/Ednrb* mutants and perhaps also our observation of normal melanocyte presence in the stria vascularis compared to past reports. In most cases, a controlled inbred strain background is a benefit in terms of maintaining phenotypic consistency. However, as described here, the phenotypic uniformity of past studies is likely to have obscured other important features of auditory maturation that are under Ednrb control. We were able to recognize these new features in the outbred background of our colony by examining a very large number of mice for all phenotypes and by segregating them through lineage-specific conditional mutagenesis. It is obvious that *EDN3* and *EDNRB* mutations in humans have variable phenotypic presentation (Pingault et al., 2010), which is a clear indication of the impact of modifier genes. Our colony resembles the human situation more accurately than any inbred background can replicate. New insights on the cell and molecular biology of auditory development that we are able to achieve with our colony may lead to new diagnostic criteria and perhaps also new therapeutic opportunities.

## Acknowledgements

No acknowledgements to provide.

## Author contributions

TM and HMS conceptualized the project and its design, JT, AD, and TM performed experiments and collected data. All authors participated in writing and editing drafts of the manuscript.

## Declaration of interests

The authors declare no competing interests.

## STAR Methods

*Key resources table:* see attached

### Resource availability

#### Lead contact

Further information and requests for resources and reagents should be directed to and will be fulfilled by the lead contact Takako Makita.

#### Materials availability

This study did not generate new unique reagents.

### Experimental model and study participant details

#### Animals

*Ednrb* (Rattner et al., 2013) (JAX:011080), *Edn3* (Baynash et al., 1994) (JAX:002516), *Wnt1Cre* (Danielian et al., 1998) (JAX:003829), *Pax2Cre* (Ohyama and Groves, 2004) (MMRRC:010569-UNC), *Phox2bCre* (Scott et al., 2011) (JAX: 016223), *Tbx18Cre* (Cai et al., 2008), *Tie2Cre* (Kisanuki et al., 2001) (JAX:008863), *ROSA26^CAG-tdTomato^* (Madisen et al., 2010) (JAX: 007914), *ROSA26^nT-nG^* (Prigge et al., 2013) (JAX: 023537), *Ednrb-EGFP* (Gong et al., 2007) (MMRRC_010620-UCD) alleles have been described previously. All experiments with animals complied with National Institutes of Health guidelines and were reviewed and approved by the Medical University of South Carolina Institutional Animal Care and Use Committee. Wild-type ICR mice used for propagation of these lines were obtained from Harlan/Envigo. All lines used in this study have been backcrossed for numerous generations to the ICR background, although because of intercrosses this was not rigorously controlled or quantified.

### Method details

#### Auditory Brain Response (ABR) and Distortion Product of Otoacoustic Emissions (DPOAE)

Postnatal day 19 mice were anesthetized with 3.0% isoflurane inhalation via a SomnoSuite small animal anesthesia system (Kent Scientific), and placed on a heating pad inside the SD1 small test enclosure (EST-LINDGREN). Needle electrodes were placed subcutaneously at the vertex (active), the ipsilateral ear (reference) and the pelvic limb (ground), and click-evoked ABR was recorded by the RZ6 signal processor (TDT) in declining 10dB steps beginning at 90dB. The ABR threshold was determined as the lowest intensity at which a recognizable wave I ABR waveform could be identified. The MF1 multi-field magnetic speaker (calibrated with BioSigRZ software (TDT) prior to each use) was used for open field ABR recording. BioSigRZ was used for ABR waveform (wave I amplitude and latency) analysis. DPOAE measurements were performed at P20 on a subset of controls and mutant mice that were found to be nonresponsive (NR) at P19. The acoustic probe containing the MF1 multi-field magnetic speakers and the EB10^+^ DPOAE microphone (TDT) was calibrated for closed field experiments and placed in the ear canal. DPOAE data were collected in response to a combination of two pure tones presented continuously at the primary frequency f1 and f2, and measured at the audiometric frequencies of 4kHz, 8kHz, 16kHz and 32kHz by the RZ6 signal processor (TDT). The primary frequency of f1 is the audiometric frequency multiplied by a factor of 0.909, and the frequency of f2 is the audiometric frequency multiplied by a factor of 1.09, and thus the frequency ratio f2/f1=1.2. Acoustic responses and DPOAE amplitudes (f_DP_=2f1-f2) with the noise floor (signal to noise ratio) were analyzed using BioSigRZ Software (TDT).

#### Cochlea preparation for histology and immunostaining

Postnatal day 20 mice were anesthetized and transcardially perfused with PBS followed by 4% paraformaldehyde. The inner ears were dissected, pierced at the apical wall of the cochlea, and postfixed in 4% PFA for 2 hr at room temperature. The entire inner ear was washed with PBS, decalcified in 8% EDTA (pH7.4) for 2-3 days, and then washed in PBS. The whole tissue was embedded in 8% low melt agarose, then vibratome sectioned at 100μm thickness.

#### Immunofluorescence staining and imaging

Inner ear sections and wholemount cochlear preparations were cryoprotected with 30% sucrose/PBS, permeabilized by three freeze-thaw cycles, and washed extensively with PBS to remove sucrose. Tissues were incubated with primary antibodies at 37°C overnight, and then with Alexa Fluor secondary antibodies (Invitrogen) for 3-5 hours with gentle agitation. Primary antibodies used in this study include BLBP (1:500; Abcam ab32423), CtBP2 (1:500; BD 612044), GFP (1:3000; Abcam ab13970), GluR2 (1:300; Millipore MAB397), HuD (1:500; SCBT sc-28299), Kir4.1 (1:400; Alomone AGP-035-GP), PECAM1/CD31 (1:200; BD 550274), Peripherin (1:200; Abcam ab99942), SYP (1:500, Abcam ab32594, 1:500, SCBT sc-17750), Tuj1 (1:1000; BioLegend MMS-435P). All immunostained sections/tissues were counterstained with DAPI, and then cleared in ScaleU2 (4M urea, 30% glycerol, 0.1% Triton-X 100). Fluorescence images were acquired using a Leica SPE confocal microscope system. To quantify the recombination efficiency (Fig. 3 and Fig. S3), five confocal images (15μm apart) were extracted from z-stack images (1μm interval) from one sagittal section containing all three turns from each cochlea analyzed, and Rosa reporter positive cells in HuD^+^ SGN neurons, BLBP^+^ satellite glia, hair cells (by anatomical location), and Kir4.1^+^ intermediate stria cells in the base through midturn were counted using the Fiji ImageJ plugin Cell Counter. To quantify IHC and OHC synaptic components (Fig. 5), three confocal z-stack images (0.5μm intervals) were acquired from base to middle turn of each cochlea and the number/volume of each fluorescence puncta was measured using the Fiji ImageJ plugin

FociPicker3D. The average of three measurements per cochlea was used for compiling data. To quantify vesicular synaptophysin in SGNs (Fig. 6a-c), five confocal images (15μm apart) were extracted from z-stack images (1μm interval) from each tissue section, and SYP^+^ area on each was measured by binary threshold selection using ImageJ normalized to the number of SGNs per image. The average of the five measurements per tissue was used for compiling data. To quantify IHC excitatory synapse components (Fig. 6d-f), five confocal z-stack images (0.5μm intervals) were acquired from base to middle turn of each cochlea and the number/volume of each fluorescence puncta was measured using the Fiji ImageJ plugin FociPicker3D. The average of five measurements per cochlea was used for compiling data.

### Statistics

All quantified data were graphed as mean±SEM, and analyzed for significance using a two-tailed Student’s t-test or ANOVA. Box and whisker plots show the mean, first and third quartiles, and full range of data except where explicitly indicated. Individually plotted outliers (Fig. 5e-f, 6h-i) were not included in the box and whisker plots.

---

**Table S1.** A genotype breakdown of control mice used for ABR study. The control group in Figure 1 is composed of animals of these specific genotypes. The 37 wild-type mice include 16 littermates of *Ednrb* mutants and 21 littermates of *Edn3* mutants.

| Genotype | n |
| --- | --- |
| *Wild-type* | 37 |
| *Ednrb*<sup>+/-</sup> | 8 |
| *Edn3*<sup>+/-</sup> | 10 |
| *Wnt1Cre/Ednrb*<sup>fl/+</sup> | 77 |
| *Pac2Cre/Ednrb*<sup>fl/+</sup> | 88 |
| *Tbx18Cre/Ednrb*<sup>fl/+</sup> | 9 |
| *Phox2bCre/Ednrb*<sup>fl/+</sup> | 12 |
| *Tie2Cre/Ednrb*<sup>fl/+</sup> | 11 |
| Total | 252 |

**Figure S1.**
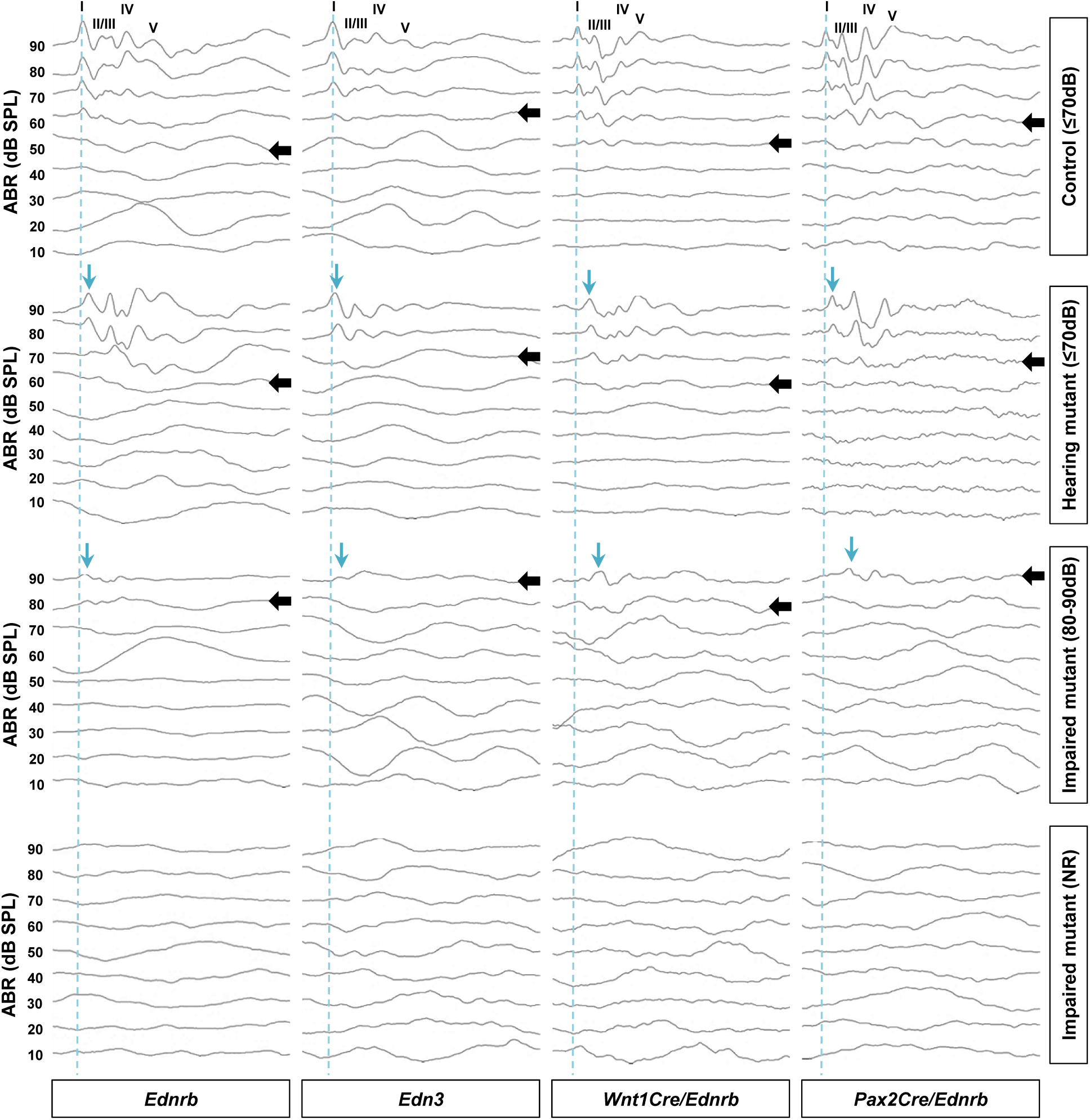
Representative ABR waveforms of P19 Edn3-Ednrb signaling mutants. Click-evoked ABR waveforms of individual animals. Because mutant mice were generated from different crosses, control mice and mutant mice with auditory phenotypes shown at right were from the same cohorts (shown at bottom) but not necessary from the same litters. The waveforms represent the responses in decreasing stimulus levels between 90dB and 10dB. Top row are the ABR waveforms of controls (see Table S1) from the indicated genetic backgrounds (i.e., littermates of mutant mice). Bottom three rows include example ABR waveforms of individual mutants from the indicated genetic background exhibiting normal hearing (ABR threshold≤70dB), impaired hearing (ABR threshold 80dB and 90dB), and total deafness (NR=no response) as indicated at the right. I-V denote the location of ABR peaks. The ABR threshold was determined as the lowest intensity at which a recognizable wave I waveform can be identified (individual threshold evaluation is noted by black arrows). Non-hearing (ABR=NR) was defined by the absence of auditory nerve response (wave I) at 90dB. Wave I represents spiral ganglion neuron activities. Blue dotted line indicates a time delay between stimulus input and SGN response in littermate control (latency). Blue arrows denote delays in wave I latency that were observed in hearing (ABR threshold≤70dB) and hearing impaired (ABR threshold 80-90dB) mutants. Compiled representations of wave I amplitude vs latency plots comparing controls vs mutants in each genetic background are shown in Fig. 1d.

**Figure S2.**
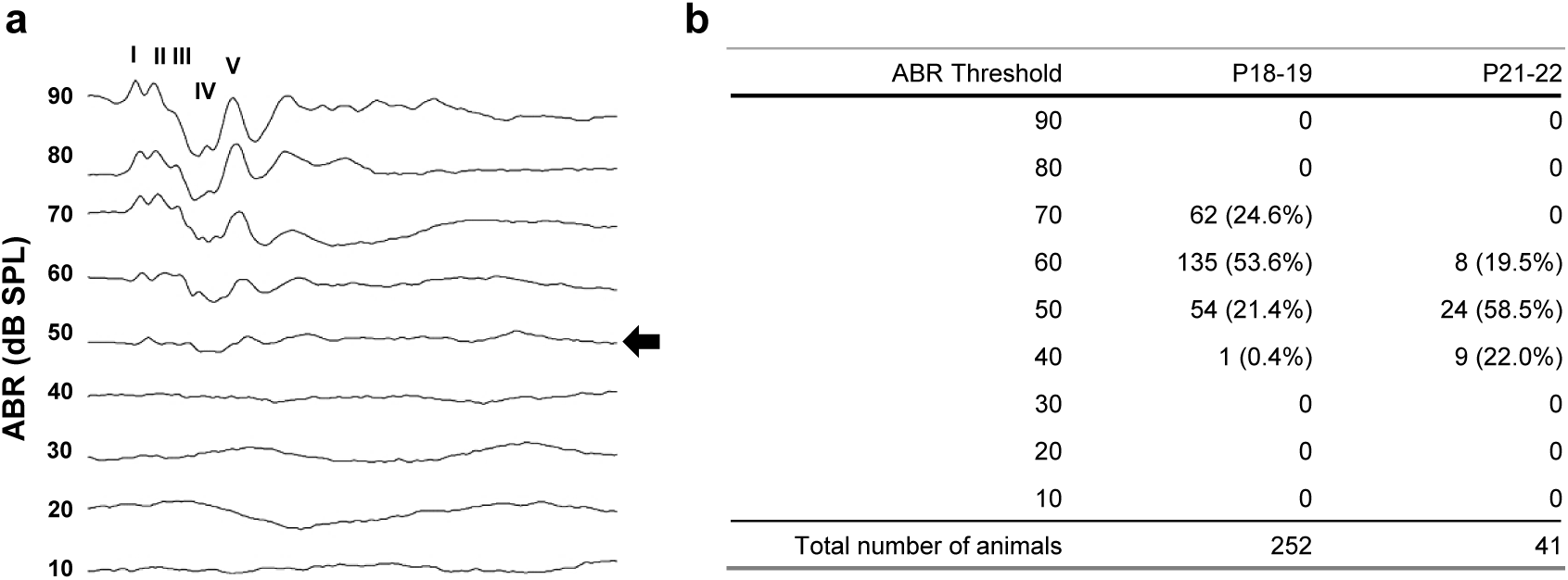
Developmental transition of hearing sensitivity in P18-19 vs. P21-22 control mice. (a) Click-evoked ABR waveforms in decreasing stimulus levels between 90dB and 10dB obtained from a P22 control mouse of ICR strain background. Black arrow point to the ABR threshold as 50dB. (b) The distribution of ABR threshold in P18-19 (n=252) vs. P21-22 (n=41) control mice. P18-19 control mice represents data sets compiled in Figure 1.

**Figure S3.**
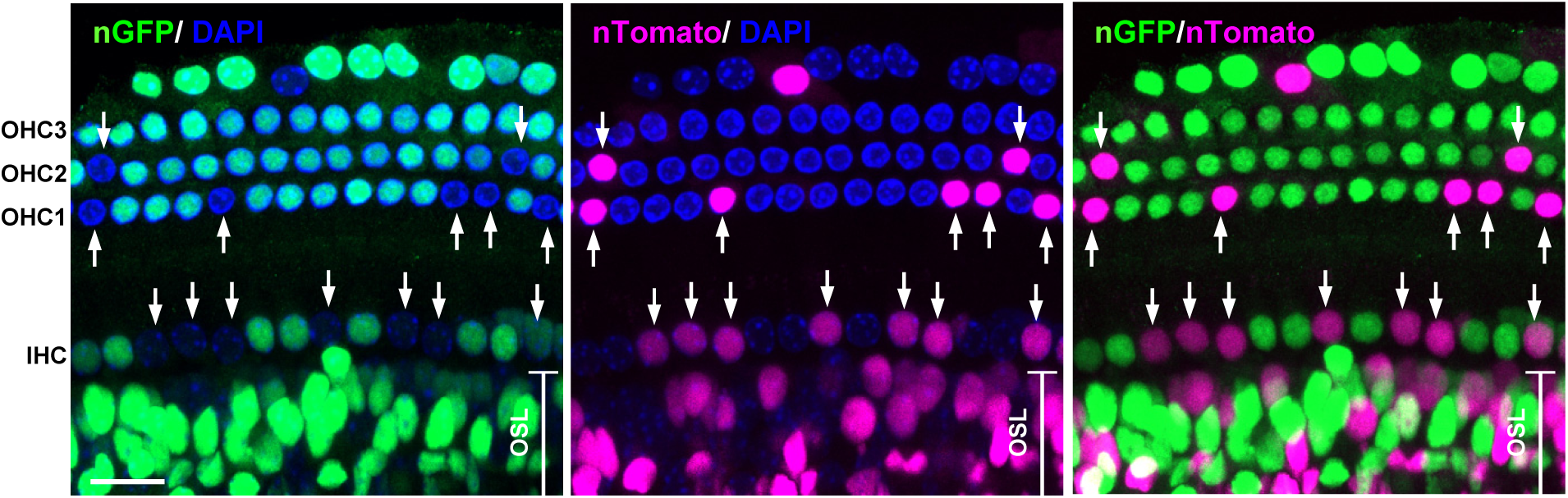
Recombination profile of *Pax2Cre* in cochlear hair cells. Wholemount cochlea preparation from P7 *Pax2Cre/R26^nT-nG^* mouse stained for GFP (green) and counterstained for DAPI (blue). Arrows point to the nuclei of non-recombined cells that express nuclear-Tomato (magenta). The osseous spiral lamina (OSL) area contains *Pax2Cre*-derived otic fibrocytes and auditory nerve associated Schwann cells which originate from non-*Pax2Cre* (*Wnt1Cre*) lineage (Fig. 3c). Scale bar; 20μm.

**Figure S4.**
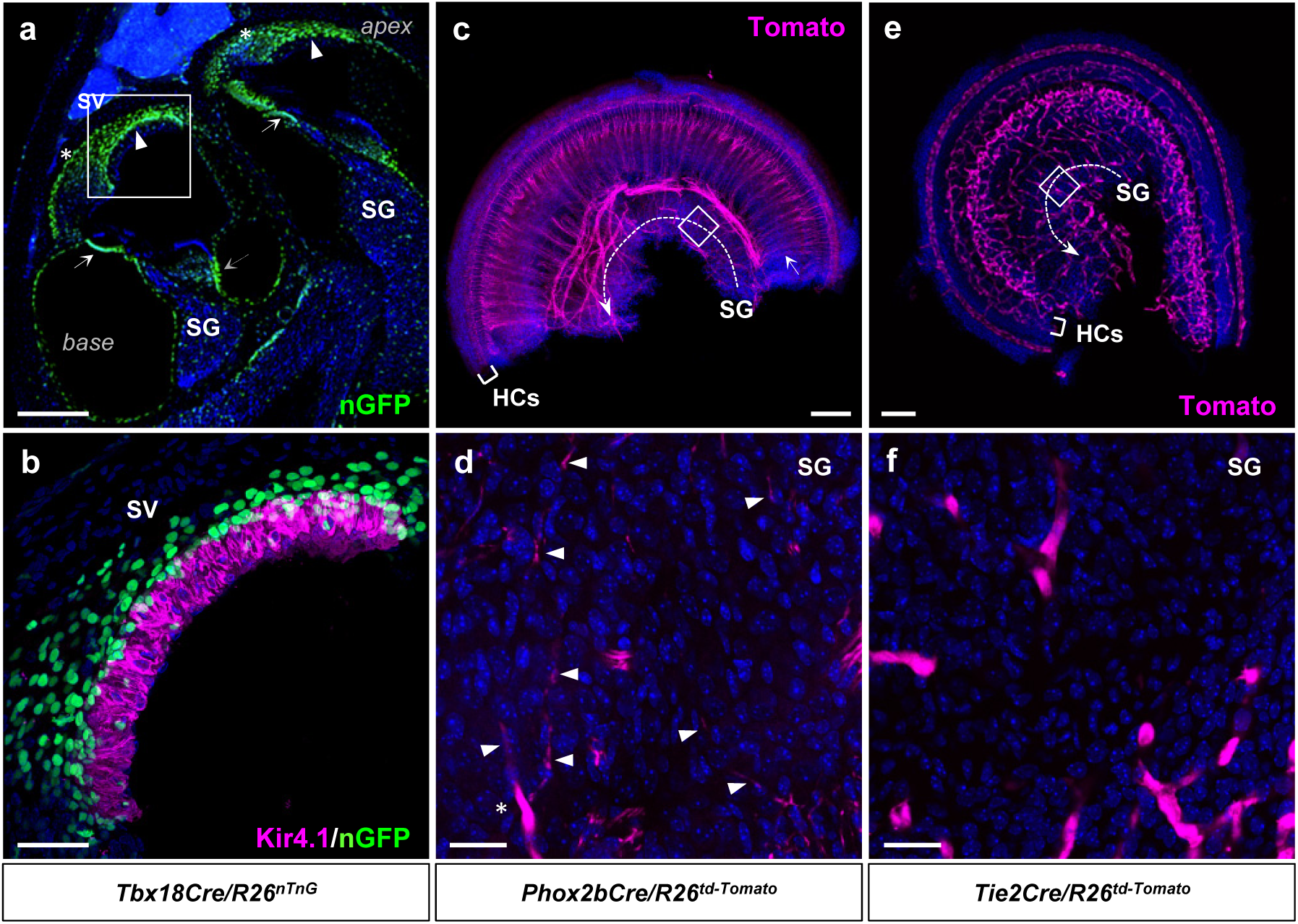
Recombination profile of *Tbx18Cre*, *Phox2bCre*, and *Tie2Cre* drivers. (**a**-**b**) Sagittal section of cochlea isolated from P19 *Tbx18Cre/R26^nT-nG^* mouse stained for GFP (green) and co-stained for Kir4.1 (magenta). (a) Low magnification view demonstrates *Tbx18Cre* activity in the stria vascularis (arrowheads) and otic mesenchyme derivatives including the basilar membrane (arrows), the lateral wall (asterisk) and the medial spiral limbus (dotted arrow). (b) High magnification of the boxed area in (a). *Tbx18Cre* labels fibrocytes in the spiral ligament and the basal stria cells which are located lateral to the Kir4.1^+^ intermediate stria cells. (c-d) Whole-mount preparation of cochlea isolated from a P0 *Phox2bCre/R26^td-Tomato^* mouse. Dotted arrow denotes the spiral ganglion (SG) and arrows point to the nerve fibers projecting towards hair cells (HCs). (d) High magnification view of the boxed area in (c). Tomato expression was not detected in neurons nor glia of the spiral ganglion, but observed in efferent nerve fibers (arrowheads) and an associated Schwann cell (asterisk) of the olivocochlear system. (e-f) Whole-mount preparation of cochlea isolated from a P0 *Tie2Cre/R26^td-Tomato^*mouse. Tomato signals represent endothelial cells of the cochlear vasculature. (f) High magnification view of the bracketed area in (e). Scale bars: 200μm (a), 100μm (b, c, e), 20μm (d, f).

**Figure S5.**
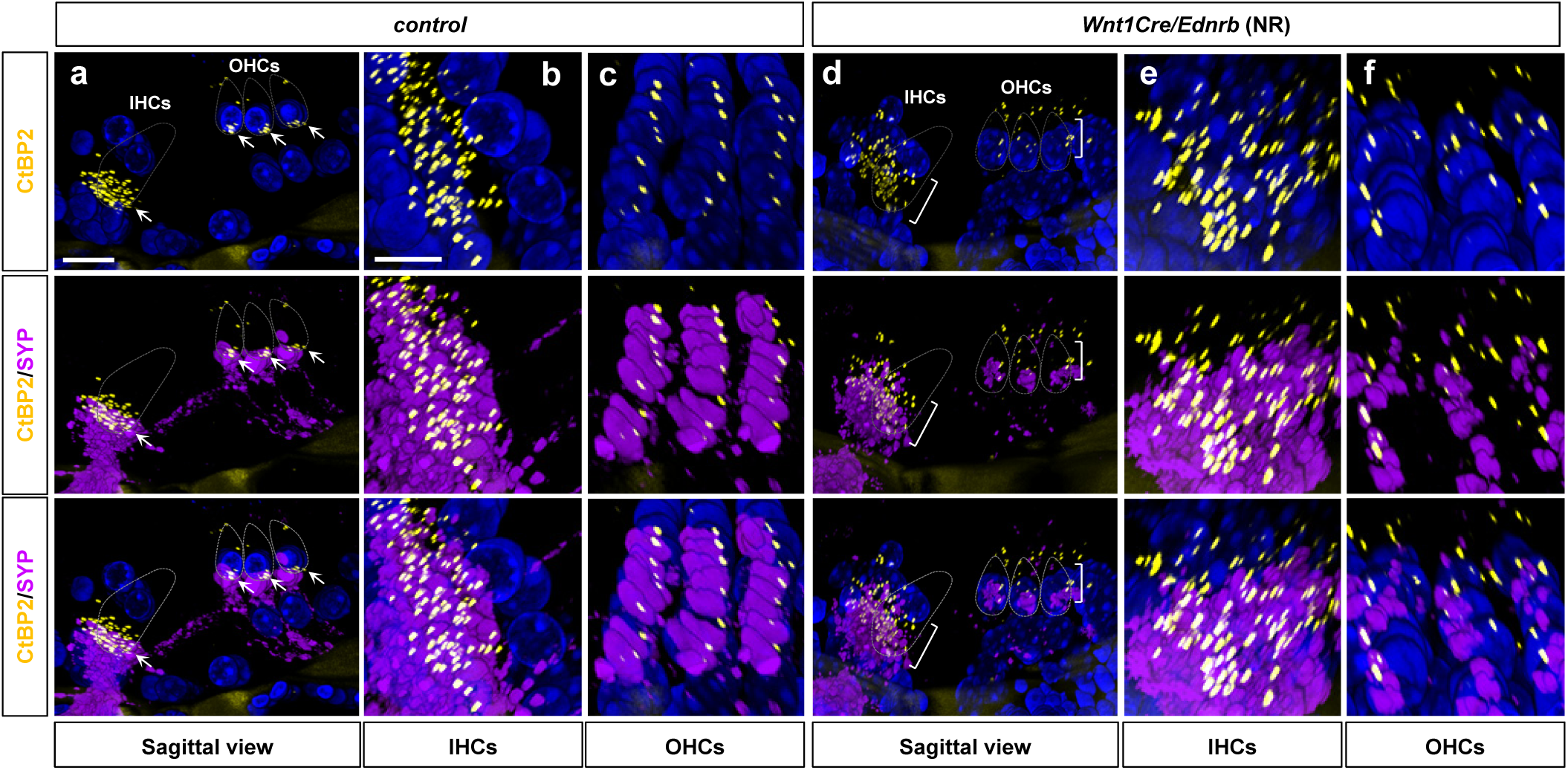
Defective pre- and post- synaptic assembly in hearing impaired *Wnt1Cre/Ednrb* mutant mice. (**a**, **d**) Confocal z-stack images from 100 μm sagittal sections of cochlea isolated from P19 deaf *Wnt1Cre/Ednrb* mutant (d) and its littermate control (a) stained for CtBP2 (yellow), co-stained for SYP (magenta) and counterstained for DAPI (blue). Dotted lines denote outlines of IHC and OHCs of the first row. (a) Arrows point to the condensed localization of CtBP2-containing synaptic ribbons alongside the SYP^+^ afferent nerve endings at the basolateral side of the control hair cells. High magnification of the rotated views of IHC and OHC synapses in (a) are shown in (b) and (c), respectively. A 100 μm section contains 8 rows of IHCs (b) and OHCs (c), and their respective afferent synapses. (d) Brackets denote scattered and disorganized localization of CtBP2 puncta within the IHC and OHCs of the first row in the *Wnt1Cre/Ednrb* mutant cochlea. High magnification of the rotated views of basolateral surface of the IHC and OHCs in (d) are shown in (e) and (f), respectively. Scale bars: 20μm (a, d), 10μm (b, c, e, f).

**Figure S6.**
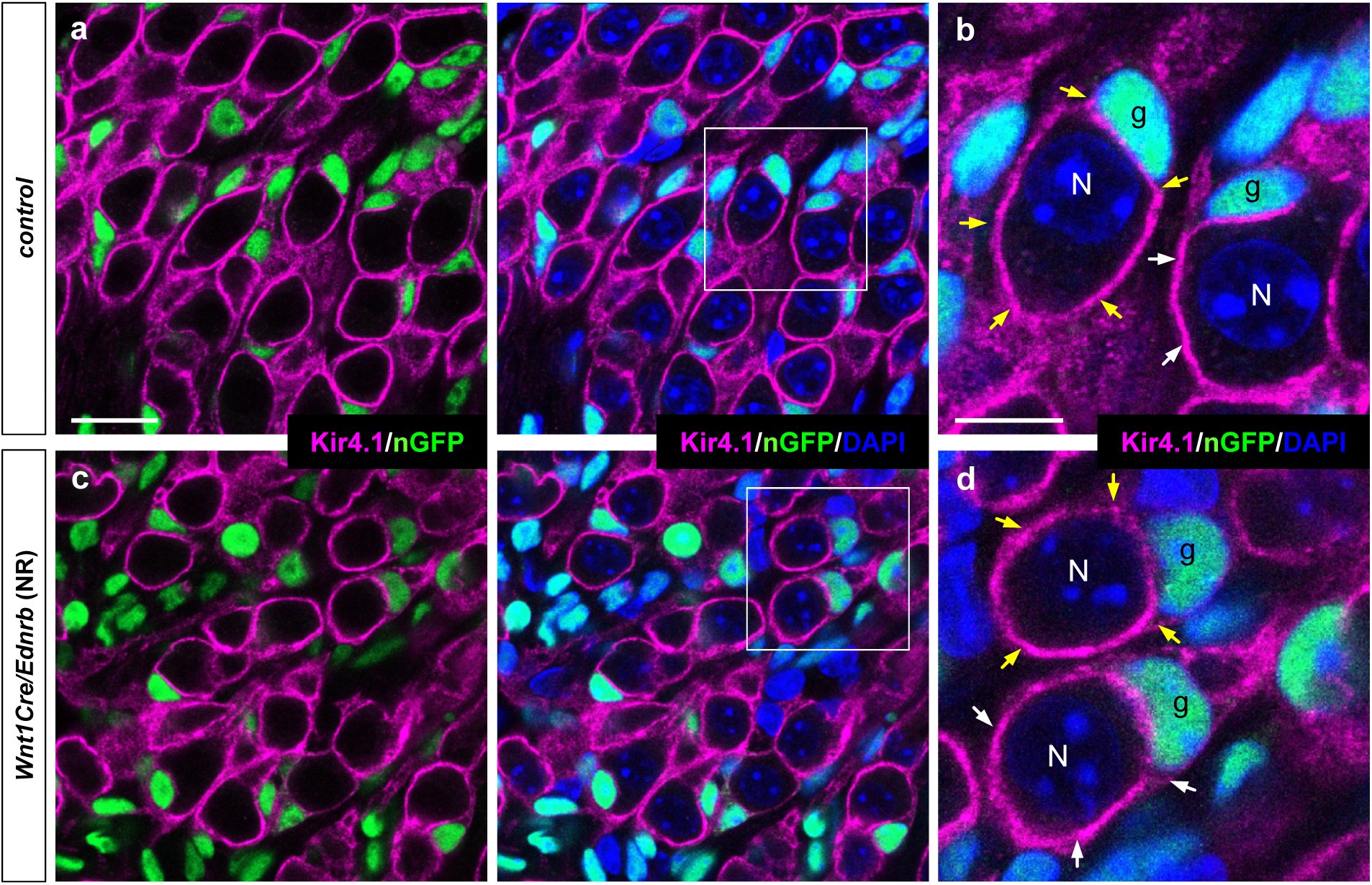
No obvious disruption in distribution and morphology of *Ednrb*-deficient satellite cells in the spiral ganglia of hearing impaired *Wnt1Cre/Ednrb* mutant mice. Confocal images of spiral ganglia isolated from P19 *Wnt1Cre/Ednrb/R26^nT-nG^* mutant (c; ABR=NR) and a littermate control (a) stained for GFP (green), co-stained for Kir4.1 (magenta), and counterstained for DAPI (blue). High magnification views of bracketed areas in (a) and (c) are shown in (b) and (d), respectively. “N” and “g” in (b) and (d) denote two individual spiral ganglion neurons and their associated satellite glia, respectively. Yellow and white arrows in (b) and (d) point to Kir4.1^+^ glial processes that ensheath their associated neurons. Scale bars: 20μm (a, c), 10μm (b, d).

**Figure S7.**
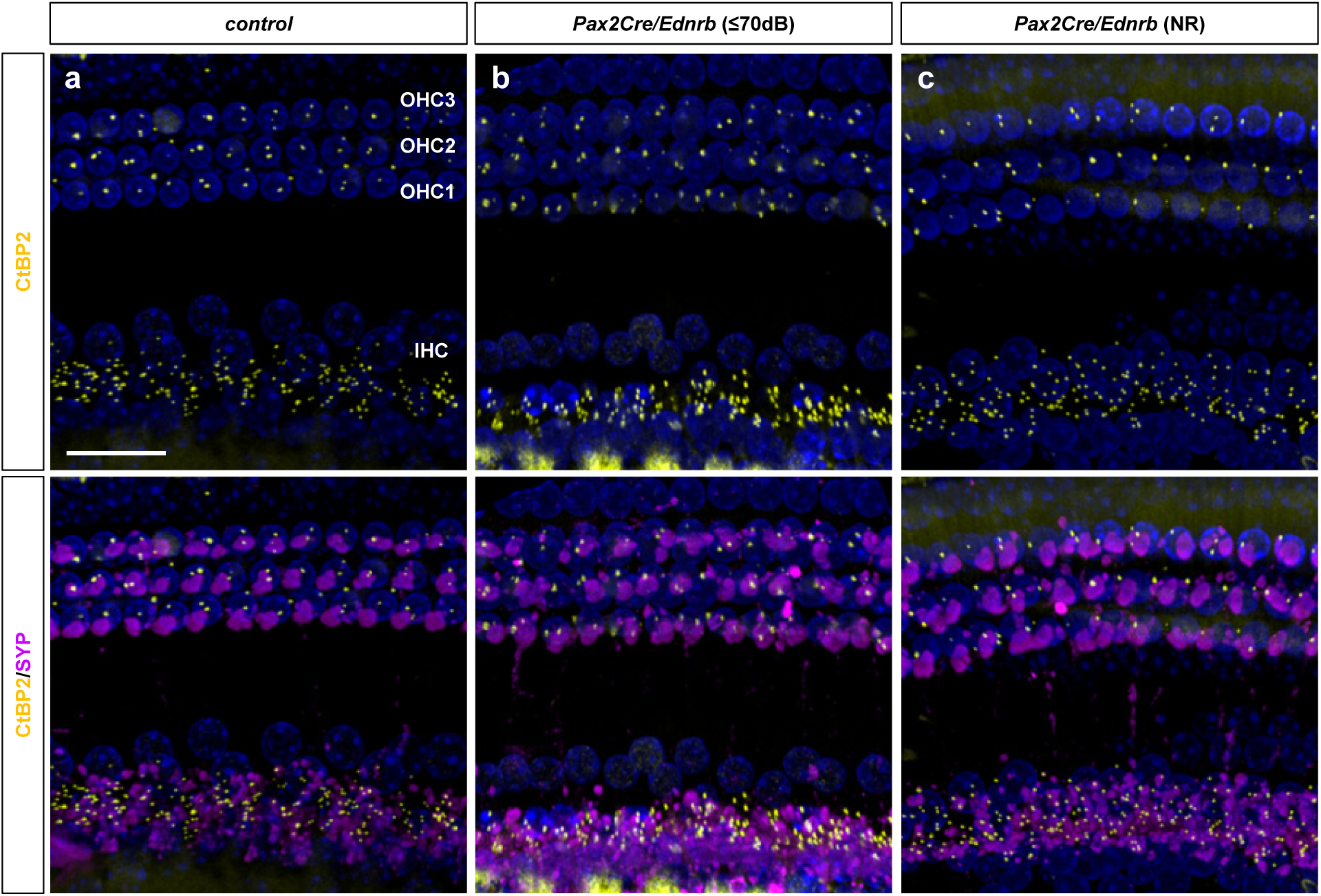
Normal pre- and postsynaptic organization in hearing impaired *Pax2Cre/Ednrb* mutant mice. (a-c) Whole-mount preparation of cochlea isolated from P19 hearing *Pax2Cre/Ednrb* (b), hearing impaired *Pax2Cre/Ednrb* (c) mutant mice and a littermate control (a) immunostained for CtBP2 (yellow), SYP (magenta) and DAPI (blue). Scale bar: 50μm (a-c).

## Notes

### Competing Interest Statement

The authors have declared no competing interest.

### Summary of Updates

Text revised; Figure 2 revised; Figure S6 revised; new figures added (Figure S2, S3, S5); author affiliations updated.

